# Pan-organ model integration of metabolic and regulatory processes in type 1 diabetes

**DOI:** 10.1101/859876

**Authors:** Marouen Ben Guebila, Ines Thiele

## Abstract

Type 1 diabetes mellitus (T1D) is a systemic disease triggered by a local autoimmune inflammatory reaction in insulin-producing cells that disrupts the glucose-insulin-glucagon system and induces organ-wide, long-term effects on glycolytic and nonglycolytic processes. Mathematical modeling of the whole-body regulatory bihormonal system has helped to identify intervention points to ensure better control of T1D but was limited to a coarse-grained representation of metabolism. To extend the depiction of T1D, we developed a whole-body model using a novel integrative modeling framework that links organ-specific regulation and metabolism. The developed framework allowed the correct prediction of disrupted metabolic processes in T1D, highlighted pathophysiological processes common with neurodegenerative disorders, and suggested calcium channel blockers as potential adjuvants for diabetes control. Additionally, the model predicted the occurrence of insulin-dependent rewiring of interorgan crosstalk. Moreover, a simulation of a population of virtual patients allowed an assessment of the impact of inter and intraindividual variability on insulin treatment and the implications for clinical outcomes. In particular, GLUT4 was suggested as a potential pharmacogenomic regulator of intraindividual insulin efficacy. Taken together, the organ-resolved, dynamic model may pave the way for a better understanding of human pathology and model-based design of precise allopathic strategies.

## Introduction

Type 1 diabetes (T1D) mellitus is a systemic disease triggered by the destruction of insulin-producing pancreatic beta cells (Maahs et al., 2010). The World Health Organization estimates that the incidence of T1D is up to 36.8 new cases per 100,000 persons per year with a 2 to 5% increase worldwide (Maahs et al., 2010). T1D remains the most prevalent type of diabetes in children and has cumbersome, lifelong effects (Maahs et al., 2010). The disease affects the production of insulin, primarily causing acute metabolic complications and coronary artery disease and resulting in high early mortality rates (Orchard et al., 2006). Additionally, the misdiagnosis of T1D in adults has been a recently acknowledged issue (Thomas et al., 2018), as a delay in implementing insulinotherapy can further exacerbate disease symptoms.

The systemic mechanism of action of insulin induces the whole-body semiology of T1D. Symptoms include several glucose-dependent metabolic processes, such as micro- and macroangiopathy, diabetic retinopathy, and nephropathy, and fatty acid-related implications, including atherosclerosis and cardiovascular disease (American Diabetes, 2010). Supplementing patients with exogenous insulin remains the gold standard treatment for T1D. Although insulin is very efficient in preventing biochemical alterations, insulin has a high within- and between-subject variability that can induce adverse reactions ranging from hypoglycemia to uncontrolled diabetes; these effects can hamper treatment compliance (Heinemann, 2002).

The wealth of data regarding T1D has helped the development of whole-body mathematical models of insulin action and disease progression. One of these models is the ordinary differential equations (ODE)-based glucose-insulin model (GIM) (Schaller et al., 2013). The GIM model includes fine-grained details regarding tissue-based insulin and glucagon action in relation to glucose dynamics, insulin-dependent receptor synthesis, and gastrointestinal hormonal regulation of glucose levels. The GIM model can reproduce the outcome of differential diagnosis tolerance tests on T1D patients (Schaller et al., 2013). The model extensively describes bihormonal regulatory events; however, the modeling of metabolism remains limited to the first step of the glycolytic pathways. The model has been used in many applications, mainly in closed-loop insulin administration in T1D patients (Schaller et al., 2016; Schaller et al., 2012; Wahdehn, 2016) and the estimation of cell proliferation rates in humans using labeled glucose (Lahoz-Beneytez et al., 2017).

Therefore, systematic analyses of disrupted metabolic processes beyond glycolysis in T1D require extended approaches. To better capture metabolism, we used an whole-body organ-resolved reconstruction of human metabolism (WBM reconstruction) (Thiele et al., 2018), which was built using the comprehensive, but generic human metabolic reconstruction (Brunk et al., 2018). The male WBM reconstruction includes the comprehensively known metabolic pathways for 20 organs, two sex organs, and six blood cells, which are anatomically accurately interconnected. Through the integration of condition-specific information, aka constraints, such as diet, gene expression, or physiological parameters, the WBM reconstruction can be converted into a personalized, condition-specific WBM model (Thiele et al., 2018). Consequently, the male WBM model enables the simulation of carbohydrate disorders and the investigation of their impact on nonglycolytic pathways. Thereby, the WBM reconstruction provides a complete picture of organ-resolved human metabolism. However, addressing the disease dynamics and patient responses to insulin requires the modeling of metabolic pathways and nonmetabolic processes as well as insulin receptor binding, signal transduction, and internalization of receptors. Recently, multiscale, multialgorithm, whole-organism dynamic models in biology, primarily models of microbiology (Bauer and Thiele, 2018; Bauer et al., 2017; Covert et al., 2008; Karr et al., 2012), plant physiology (Grafahrend-Belau et al., 2013), human physiology and xenobiotic metabolism (Cordes et al., 2018; Guebila and Thiele, 2016; Krauss et al., 2012; Thiele et al., 2017; Toroghi et al., 2016), have increased in complexity.

Accordingly, we coupled the male WBM reconstruction with the dynamic ODE-based GIM model. The multiscale, hybrid model (dWBM) enabled the representation of T1D through both glycolytic target effects and off-target effects that were mapped onto the dWBM model using gene expression data. In particular, the effects of insulin were considered by including an additional dynamical model representing the insulin impact on liver metabolism. The resulting model included the target and off-target effects of both T1D and insulin, and i) enabled the quantification of disrupted metabolic processes in T1D in relation to the disease symptomatology, ii) allowed an assessment of the insulin nonglycolytic effects, and iii) shed new light on the mechanisms underlying the inter- and intra-individual variability in insulin effects. The hybrid dWBM model provides a basis for the organ-specific integration of biological data and paves the way for predictive human physiology.

## Results

Our approach to constructing the whole-body model of carbohydrate metabolism consisted of integrating the dynamical regulatory model of glucose-insulin metabolism with a male WBM model. The resulting hybrid model (dWBM) had a better predictive ability than each model alone and showed notable differences between healthy and T1D metabolic states. The off-target effects of both T1D and the insulin treatment were modeled at the metabolic level using gene expression data of T1D and the metabolomics of insulin treatment. Therefore, hybrid model included glucose metabolism at different scales (gene expression, regulation loops, and metabolism). The modeling of both the target and off-target effects of T1D and insulin allowed us to address the between- and within-patient variability in response to insulin.

### Whole-body flux dynamics discriminate between healthy and type 1 diabetes states and insulin and glucose challenges

In the clinical routine, the diagnosis of diabetes mellitus is confirmed by antibody detection and tolerance tests (Atkinson et al., 2014). To identify the global metabolic shift induced by T1D and glucose or insulin challenges during the tolerance tests, we built a hybrid model (Figure 1) including continuous and discrete dynamics. We then assessed the effects of the different tolerance tests in healthy (Figure 2-A) and T1D (Figure 2-B) models on whole-body, organ-wide metabolic fluxes (Methods-Simulation setting). A support vector machine (SVM) classifier segregated the simulation results of the insulin tests, i.e., intravenous insulin tolerance test, subcutaneous insulin bolus, and subcutaneous insulin infusion, as well as the glucose tests, i.e., intravenous glucose tolerance test (IVGTT), liquid oral glucose solution, and solid meal (Figure 2-C), at five and 15 minutes after perturbation and at all five-minute intervals of the 600-minute simulation, which were aggregated in the binary classifier (Figure S2, AUROC=0.82). Through the application of gene expression data from the T1D human pancreas and pancreatic islets (Planas et al., 2010) (Table S1) as additional constraints on the pancreas in the dWBM enabled to account for metabolic effects that were not directly linked to the decrease in insulin levels and the consequent increase in glucose levels represented by GIM. Expectedly, the off-target effects were primarily represented in inflammation and immune system disorders (Figure S1-A-B), and the 24 metabolic genes that were obtained through the intersection of genes in WBM and differentially expressed genes in T1D (Figure S3) represented the main features of T1D, e.g., disruption of glucose transport and gluconeogenesis (Figure S1-C-D-E).

**Figure 1.**
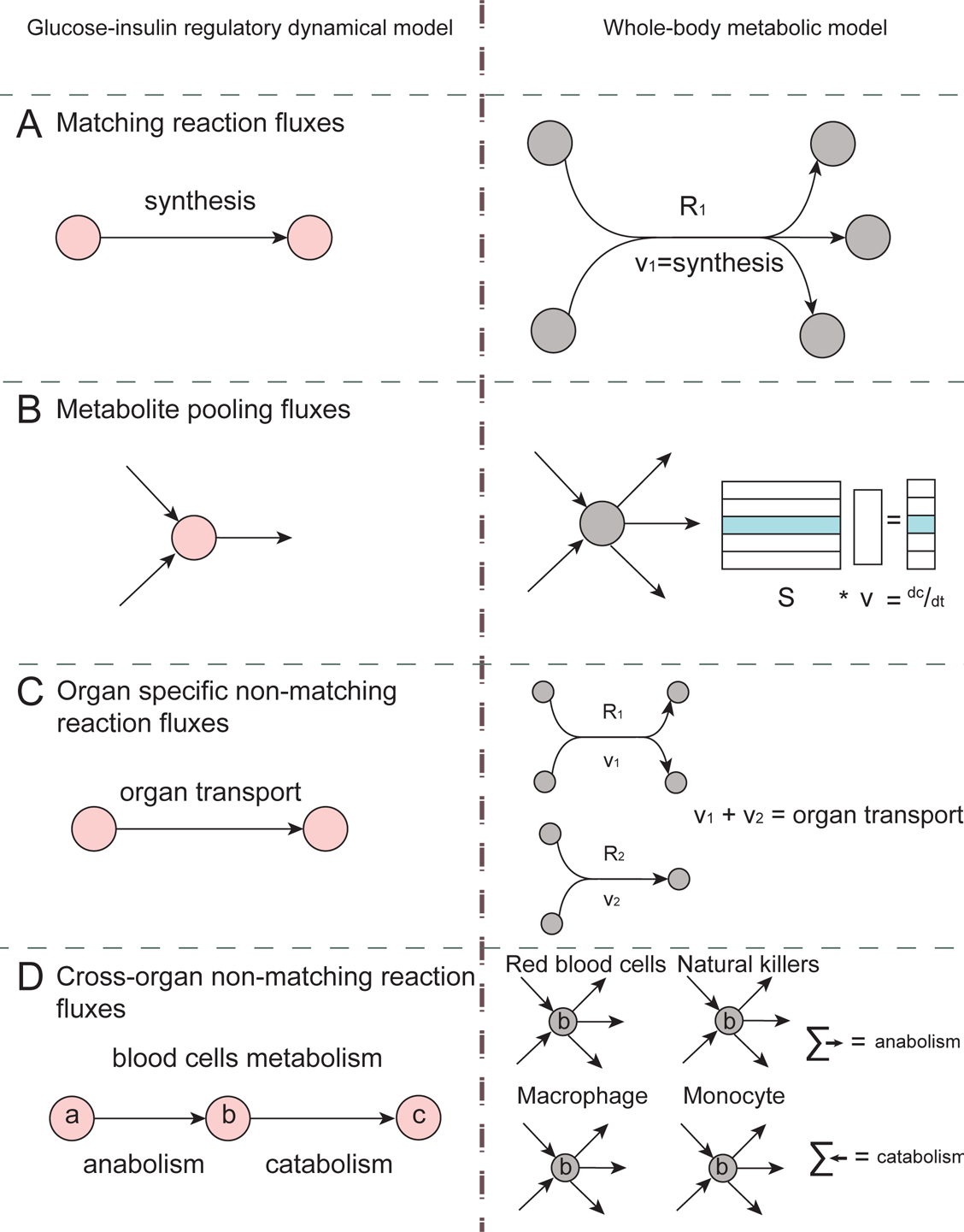
Applied constraints from the GIM model to the dWBM model. The constraints followed the following four main scenarios: **A**- Matching reaction fluxes correspond to the case, in which a reaction in the GIM model is represented in the same way as in the WBM model. In this case, the fluxes fully match. **B**- Metabolite pooling fluxes correspond to the case, in which the rate of change in a metabolite in the GIM model is set as a constraint in the right-hand side (*b* vector) of the linear programming problem in the WBM model. **C**- Organ-specific nonmatching reaction fluxes correspond to the case in which one reaction in the GIM model corresponds to more than one reaction in the same organ in the WBM model. **D**- Cross-organ nonmatching reaction fluxes correspond to the case, in which one tissue in the GIM model is represented by at least one compartment in the WBM model. The sum of the cross-organ anabolic (catabolic) fluxes is set equal to the anabolic (catabolic) fluxes in the GIM model. This case is typically related to blood cells.

**Figure 2.**
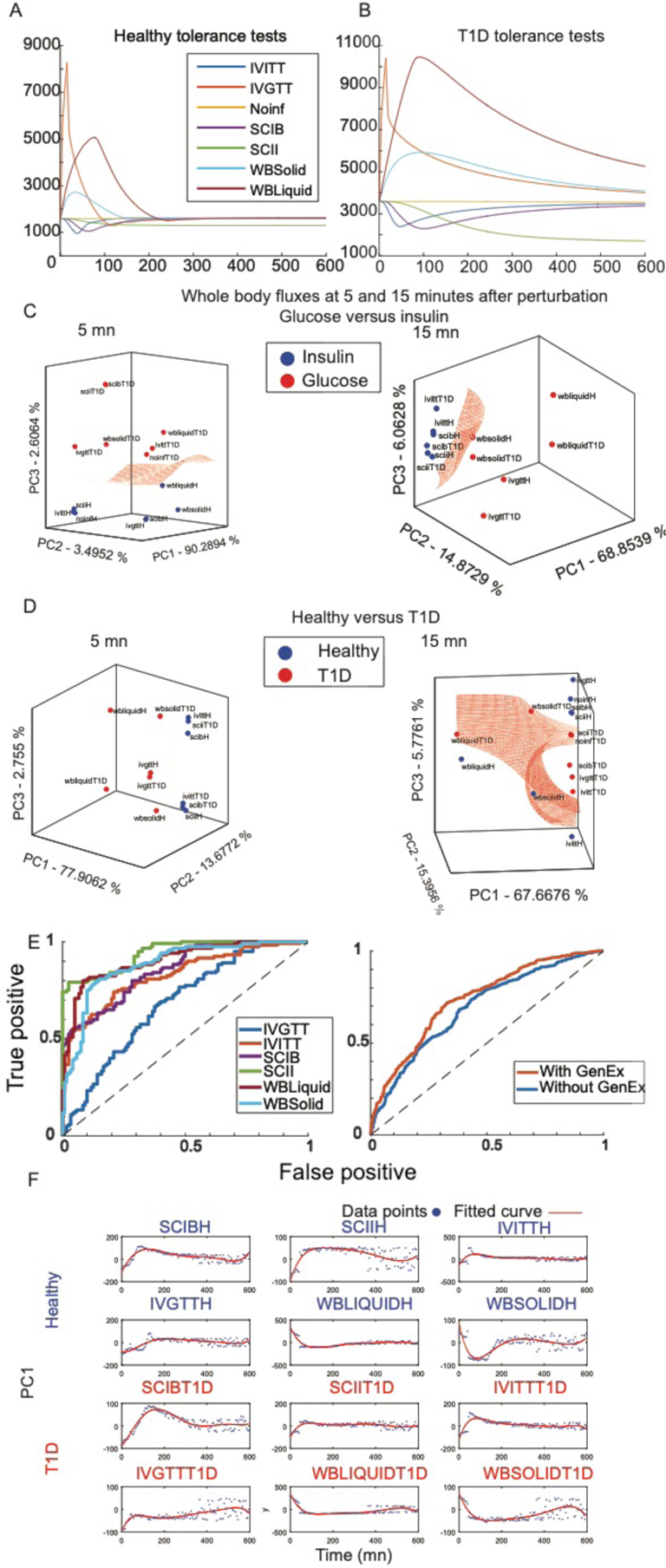
Whole-body reaction flux dynamics classify T1D and healthy states. **A**- Time course of glucose in peripheral blood in healthy and **B**- T1D states in different tolerance tests using the GIM model. **C**- Principal component analysis of whole-body metabolic fluxes in insulin and glucose challenges and **D**- in healthy and T1D dWBM models at five and 15 minutes after the perturbation using the first three components. The SVM Gaussian boundary was plotted to assess the separation among the classes. **E**- ROC curves of T1D classification. Whole-body flux dynamics over the 600-minute simulation discriminate between healthy and T1D states in each tolerance test alone (left) and in a combination of all tolerance tests in a binary classifier (right), where adding gene expression constraints results in a higher T1D predictive capability of the model. **F**- Time course of the first principal component (PC1) of the flux vector in each time step in different tolerance tests showed a shift in global metabolism as a result of glucose dynamics. IVITT: intravenous insulin tolerance test, IVGTT: intravenous glucose tolerance test, Noinf: baseline glucose concentration, SCIB: subcutaneous insulin bolus, SCII: subcutaneous insulin infusion, WBSolid: solid meal, WBLiquid: oral liquid glucose solution.

The gene expression-derived constraints represented both glycolytic effects and the immune system disruption effect on metabolism, which allowed the segregation of the whole-body flux in the healthy and disease dWBM models (Figure 2-D) at five and 15 minutes after perturbation and throughout the entire duration of the simulation (Figure 2-E, left panel, AUROC>0.68). Particularly, the classification of the glucose and insulin challenges improved (Figure S2) when the gene expression constraints were applied to the pancreas in the dWBM models (AUROC=0.82 and AUROC=0.8, respectively). Furthermore, compared to the dWBM models that did not have these applied constraints (AUROC=0.69), the classification of T1D improved (Figure 2-E (right panel), AUROC=0.73), highlighting the importance of further disease specific constraints.

The global metabolic change across the dWBM models was further supported by the time course of the first component (PC1) in the different tolerance tests (Figure 2-F). The addition of the off-target effects of T1D using gene expression data of the pancreas enabled the accurate classification of the T1D and healthy dWBM models in the whole-body flux space. We conclude that the simulations with the dWBM model on a whole-body scale were predictive of both the condition, i.e., T1D, and challenge, i.e., insulin and glucose, and allowed the assessment of disrupted metabolic pathways in T1D.

### Differential reaction fluxes between healthy individuals and type 1 diabetes patients

The T1D and healthy dWBM models were employed to assess the effect of T1D on whole-body metabolism. Using flux variability analysis (FVA) (Mahadevan and Schilling, 2003), we compared the flux span, which refers to the difference between the maximal and minimal possible flux through each reaction in the WBM and dWBM models. As expected, the dWBM model had a reduced steady-state solution space (Figure 3-A) compared to the WBM model. This result means that the set of obtained flux solutions was further constrained to biologically relevant phenotypes with respect to glucose metabolism. The T1D dWBM model showed a smaller flux span than the dWBM healthy model (Figure 3-A) and thus, a decreased metabolic flexibility towards the glycolytic disruption that characterize the disease pathophysiology.

**Figure 3.**
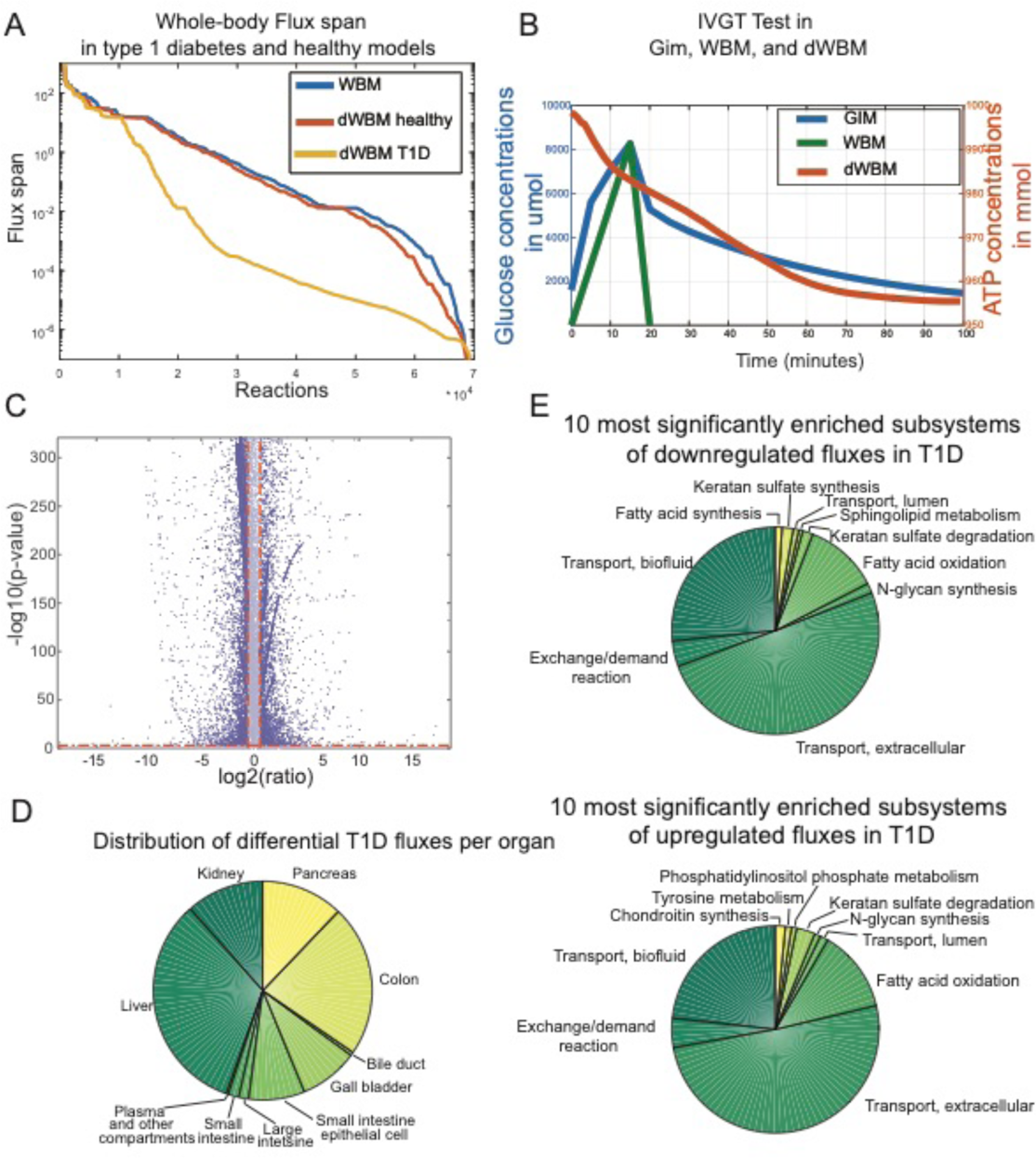
Multiscale whole-body model identified disrupted metabolic processes in type 1 diabetes mellitus. **A**- Comparison of flux variability analysis of the pancreas between the WBM and dWBM models in healthy and T1D state. **B**- Time course of glucose in peripheral blood in IVGTT modeled by the WBM model and the GIM model and theoretical amount of ATP in adipocytes during IVGTT as predicted by the dWBM model. **C**- Volcano plot of the differential reaction fluxes (ratio of fold change) in the healthy and T1D models. **D**- Differential flux distribution by organ and **E**- subsystem (p<0.05).

We compared the predictive capabilities of the WBM, GIM, and dWBM models with respect to metabolite dynamics in an IVGTT setup (see Methods). While the GIM model predicted the glucose kinetics with top-down estimated parameters based on *in vivo* measured concentrations (Schaller et al., 2013), the WBM model failed, as to be expected, to predict the glucose dynamics because of the lack of insulin and glucagon regulation (Figure 3-B). The dWBM model performed equally well to the GIM model and could predict the dynamics of the metabolites involved in the reactions that allow metabolite accumulation and depletion (i.e., reactions containing metabolites that are not at a steady state), such as the ATP demand in adipocytes during IVGTT.

Additionally, the baseline glucose levels in the healthy and T1D dWBM models could inform about steady-state glucose fluxes as illustrated by the fold change in the reaction fluxes between the healthy and T1D states (Figure 3-C). FVA was carried out on the healthy and T1D irreversible dWBM models, in which all reversible reactions were replaced by corresponding forward and reverse reactions, to guarantee positive flux values for the subsequent fold change analyses. Using the FVA results data, we compared flux distributions rather than single flux values (see Methods-differential reaction fluxes between healthy and T1D models) and performed a fold change analysis of the flux distributions. The obtained volcano plot showed significantly different reaction fluxes under both conditions. After mapping the irreversible reactions to their original identifiers, 33,526 reactions of all 80,016 reactions (42%) exhibited a change in the flux value between the conditions, including 6,602 reactions (8%) that were significantly increased and 15,205 reactions (19%) that were significantly decreased above 50% (fold change greater than 1.5) of their healthy values (p<0.001, t-test). To find small molecules that could potentially reverse the metabolic signature of T1D, we queried the LINCS database (Duan et al., 2014) for the genes associated with increased and decreased reaction flux values. Among the small molecules found to reverse the expression of those genes, Mibefradil and Amlodipine had very good gene coverage (Table S6). Additionally, the enrichment of the differentially active reactions in the metabolic subsystems (Figure 3-D) and organs (Figure 3-E) showed an expected systematic and ubiquitous deregulation of glucose metabolism, particularly in the target organs (i.e., pancreas, liver, and kidney).

The subsystem responsible for transport reactions was the most affected since it covered the regulation of glucose exchange from producing organs, e.g., from glycogenolysis in the liver to energy-requiring processes. Overall, the differential fluxes of the reactions were related to the known semiology of T1D in different organs (Table S4), e.g., cardiovascular disease, retinopathy, ketoacidosis, and liver glycogen deposition. Notably, the disruption of the transport system in T1D (Figure 3-E) suggested that insulin played a prominent role in maintaining global interorgan communication.

### Prediction of exogenous insulin off-target effects

The physiological effects of insulin mainly affect the uptake of glucose by organs. A recent study (Yugi et al., 2014) has suggested that insulin regulates the liver isoform of phosphofructokinase (PFK) such that upstream fluxes of PFK are downregulated and downstream fluxes are upregulated. Therefore, we used the dWBM model to simulate a subcutaneous insulin bolus and then to predict flux patterns with regards to insulin regulation. The liver glycolysis fluxes (Figure S6) were predicted correctly in three out of eight regulatory flux patterns, while one flux remained invariant, and four fluxes were predicted incorrectly. As the aforementioned regulation was not captured in the dWBM model, we used an additional dynamical model (Yugi et al., 2014) to represent the off-target effects of insulin regulation in the liver. This ODE model was coupled to the dWBM model via the concentrations of eight liver metabolites (fructose-6-phosphate, fructose-1,6-diphosphate, phosphoenolpyruvate, isocitrate, 2-oxoglutarate, malate, fructose-2,6-biphosphate, and citrate), which captured both the T1D and insulin target and off-target effects. In addition to glycolysis, insulin influences amino acid uptake, particularly, the uptake of large and neutral amino acids (LNAA), resulting in a decrease in blood concentrations of LNAAs (Lipsett et al., 1973). The resulting diWBM model predicted a decrease in all LNAAs in the blood (Figure S6), 60 minutes post-insulin administration after a transient increase at 30 minutes.

To infer the metabolic effects of insulin, interpreting individual flux solutions of the diWBM model is inefficient given the high dimensionality of the model and the existence of alternate optimal solutions (AOS) (Mahadevan and Schilling, 2003; Thiele et al., 2010). To characterize the AOS space, FVA provides an empirical flux probability density per reaction. We compared the probability density estimates of the flux values in T1D with and without insulin administration. As expected, glucose uptake increased in the muscle, adipocytes, and lungs but remained unchanged in the liver and brain (Figure 4) where insulin-induced glucose receptors are absent in the GIM model. In fact, the vital roles of the brain and liver necessitate a constant supply of glucose independently from external factors (Hasselbalch et al., 1999; Vella et al., 2002). Insulin stimulated the activity of glycogen synthase and hexokinase (Figure 4), contributing to its overall anabolic effect. The activity of glucose-6-phosphatase increased, and the activity of phosphofructokinase decreased, which is inconsistent with empirical observations, suggesting that insulin regulates the two enzymes by nonmetabolic mechanisms. The uptake of phosphate by adipocytes also increased, which is related to its known depletion after continuous insulin injection (Liamis et al., 2014), while the uptake of potassium did not significantly change. In relation to fatty acid biosynthesis, insulin enhanced the synthesis of lipoproteins in the liver and inhibited the oxidation of fatty acids and diacyl glycerol lipase in adipocytes. The prediction of the triglyceride levels (Figure S7) showed an increase in adipocytes following the administration of insulin. Finally, the uptake of glutamine in red blood cells and serine in the spleen (Figure 4) supported our earlier findings (Figure S6) regarding the lower uptake of amino acids and the subsequent increase in blood concentrations immediately after the administration of insulin.

**Figure 4.**
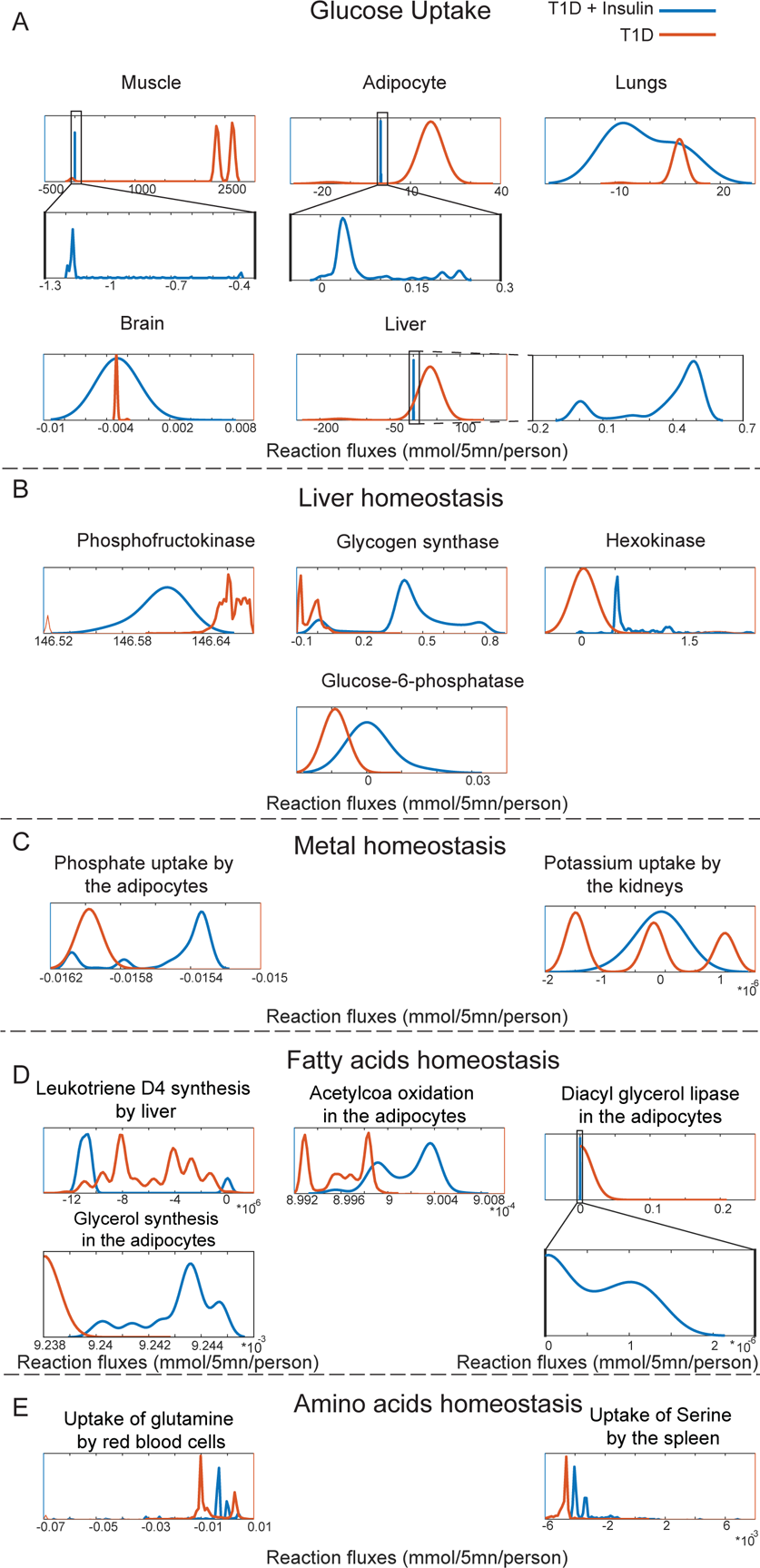
Insulin metabolic off-target effects assessed by the probability density estimates of the reaction flux values. The fluxes include **A**- glucose uptake, **B**- liver glycolytic reactions, **C**- metal homeostasis, **D**- fatty acids reactions, and **E**- amino acid uptake in various organs after a subcutaneous insulin administration.

We also investigated the anabolic effect of insulin on interorgan crosstalk in the T1D diWBM model, with 236 metabolites being exchanged between the muscle, kidney, brain, liver, adipocytes, and pancreas. In comparison, after insulin administration in the T1D diWBM model, an increase in the set of exchanged metabolites to 292 metabolites was found between the same organs (Figure S8).

### Between-subject variability is reflected in the citric acid cycle and oxidative phosphorylation

We now used the diWBM model to investigated the effects of within- and between-patient response to insulin administration and potential mechanisms underlying the insulin response interindividual variability. The set of kinetic parameters of the GIM model with differential values between T1D and healthy individuals (Table 1) (Schaller et al., 2013) was varied (Figure 5-A) to reproduce the clinically observed 25-35% (Heinemann, 2002) interindividual variability in 31 synthetic T1D patients (30 simulated patients and 1 average reference patient) (Figure 5-B). The obtained between-subject variability was 30.11%, which is consistent with the empirical values (Heinemann, 2002). For each synthetic patient, the personalized diWBM was simulated for 600 minutes after a subcutaneous injection of insulin bolus.

**Table 1.** Selected whole-body parameters in inter-individual variability simulations.

| Parameter | Parameter | Description |
| --- | --- | --- |
| P_1711 | Liver\|Interstitial\|Glucagon_0 | Baseline glucagon level in the liver |
| P_2049 | Pancreas\|Interstitial\|Insulin_0 | Baseline insulin level in the pancreas |
| P_322 | Fat\|Endosome\|End_IR_0 | Adipocyte baseline insulin receptor in the endosome |
| P_1757/P_1755 | Liver\|Intracellular\|End_IR_0_liv | Liver baseline insulin receptor in the cell |
| P_1978 | Muscle\|Endosome\|End_IR_0 | Muscle baseline insulin receptor in the endosome |
| P_323 | Fat\|Endosome\|End_IRp_0 | Adipocyte baseline insulin phosphorylated receptor in the endosome |
| P_1760 | Liver\|Intracellular\|End_IRp_0_liv | Liver baseline insulin phosphorylated receptor in the cell |
| P_1979 | Muscle\|Endosome\|End_IRp_0 | Muscle baseline insulin phosphorylated receptor in the endosome |
| P_293 | Fat\|Intracellular\|IR_cell_0 | Total concentration of adipocyte intracellular insulin receptor |
| P_1753 | Liver\|Intracellular\|IR_cell_0 | Total concentration of liver intracellular insulin receptor |
| P_1950 | Muscle\|Intracellular\|IR_cell_0 | Total concentration of muscle intracellular insulin receptor |
| P_294 | Fat\|Intracellular\|IR_p_0 | Adipocyte baseline insulin phosphorylated receptor in the cell |
| P_1758/P_1727 | Liver\|Intracellular\|IR_p_0 | Liver baseline insulin phosphorylated receptor in the cell |
| P_1949 | Muscle\|Intracellular\|IR_p_0 | Muscle baseline insulin phosphorylated receptor in the cell |
| P_1814 | Liver\|ReceptorRecyclingFactor-End | Liver receptor recycling factor in the endosome |
| P_1994 | Muscle\|ReceptorRecyclingFactor | Muscle receptor recycling factor |
| P_338 | Fat\|ReceptorRecyclingFactor | Adipocyte receptor recycling factor |
| P_1811 | Liver\|ReceptorRecyclingFactor | Liver receptor recycling factor |

**Figure 5.**
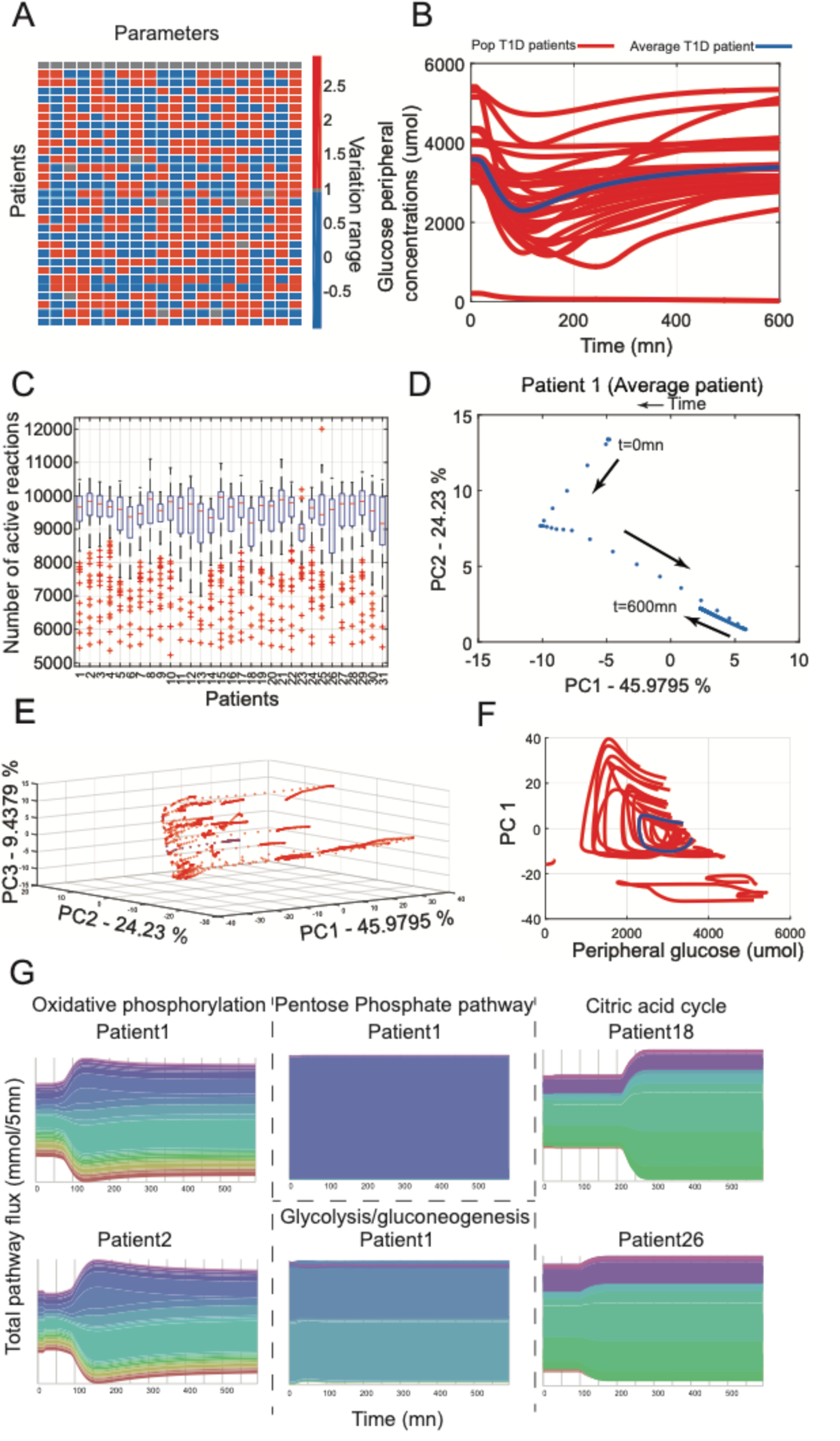
Interindividual variability in the insulin response is reflected in key pathways of metabolism. **A**- Varying a set of individual parameters within the reported interindividual variability range in the dynamical model yields **B**- different peripheral glucose concentrations in response to the insulin injection. **C**- Distribution of the active reactions over the simulation time per patient. **D**- Evolution of the metabolic fluxes as represented by PCA over the simulation time in an average patient (patient 1) and **E**- a population of synthetic patients. **F**- Insulin-induced hysteresis between glucose peripheral blood levels and whole-body metabolism. **G**- Citric acid cycle and oxidative phosphorylation total flux reflect between- patient variability, while glycolysis and the pentose phosphate pathway are stable following the perturbation. The flow chart was created using RAWGraphs (Mauri et al., 2017).

The distribution of active reactions, i.e., reactions with a nonzero flux, across patients and across time was very similar (Figure 5-C, Figure S5), which showed that the between-patient differences were not reflected in the network topology. Reducing the dimensionality of the flux distributions through Principal Component Analysis (PCA) (Figure 5-D) showed that it was rather the changes in reaction flux values across time following the administration of insulin that were differential between patients. Particularly, the final metabolic state differed from the initial state, confirming that the metabolic change was due to an action of insulin (Figure 5-D). Moreover, each individual had a differential metabolic shift, which was reflective of the quantitative variation in the flux values across pathways (Figure 5-E). Interestingly, we observed that insulin exerted a hysteresis between the peripheral glucose concentrations and whole-body metabolism (Figure 5-F) that was achieved differently in the virtual population.

We further investigated the impact of interindividual variability on different metabolic pathways. In particular, the pentose phosphate pathway and glycolysis were remarkably resilient against the insulin-induced perturbation, while the citric acid cycle and oxidative phosphorylation total whole-body flux over time showed patient-specific variation (Figure 5-G). Taken together, interindividual variability to insulin was mainly reflected in the flux values of the reactions in the citric acid cycle and oxidative phosphorylation pathways.

### Within-subject variability in the insulin response

Within-subject variability in the insulin response results from internal, endogenous factors that are specific to the particular metabolic state of the individual, e.g., the postprandial state and physical activity. To determine the influence of the metabolic state on the peripheral glucose dynamics, we set the kinetic parameters of the simulated patient to the reported population average in the GIM model and varied the metabolic reactions in the diWBM model. For each metabolic state, we generated random objective coefficient weights for each reaction in the diWBM model (Figure 6-A). The weights could be lower, higher, or equal to the average profile. In the latter case, the simulated profile approached the predictions of the GIM model. In each set of internal perturbations (input) to metabolism, i.e., increase or decrease in reaction weights, we calculated the minimum glucose concentration and the final concentration ten hours after a subcutaneous insulin injection simulation (output) (Sarkar and Sobie, 2010) using direct coupling (Figure S4, Figure 1). In all cases, the time interval was decreased from five minutes to 2.5 minutes as both integration failures and conflicting constraints emerged with higher time intervals (Methods). The reactions of the diWBM model fell into two classes, i.e., reactions unique to the WBM model, and reactions shared by both the GIM and the WBM models, which are referred to as interface reactions (Wahdehn, 2016). Only changing the coefficients of the interface reactions and simulating the diWBM models with flux balance analysis (FBA) for ten metabolic states resulted in a smooth set of glucose dynamics (Figure 6-B). To obtain a complete view of the within-patient variability, the reactions unique to the WBM model were also included. Using FBA to simulate ten metabolic states of the diWBM model, the glucose profiles were nonsmooth profiles with nonbiological concentrations or the profiles terminated during the simulation (Figure 6-C). To ensure the smoothness of the system and minimal debugging to resolve conflicting constraints, we devised a simulation framework (CRONICS) that includes additional constraints to account for the dependencies between time steps (Figure S4). The simulated profiles showed smooth concentrations profiles with models including both dynamical constraints and whole-body fluxes (Figure 6-D). Only two simulations permanently failed because they were depleted of glucose and were removed from the subsequent analysis. The 29 simulated metabolic states with the final setting resulted from the perturbation of 2,817 reactions. Each metabolic state consisted of a set of perturbed reaction activities as depicted by the random coefficient matrix (Figure 6-E).

**Figure 6.**
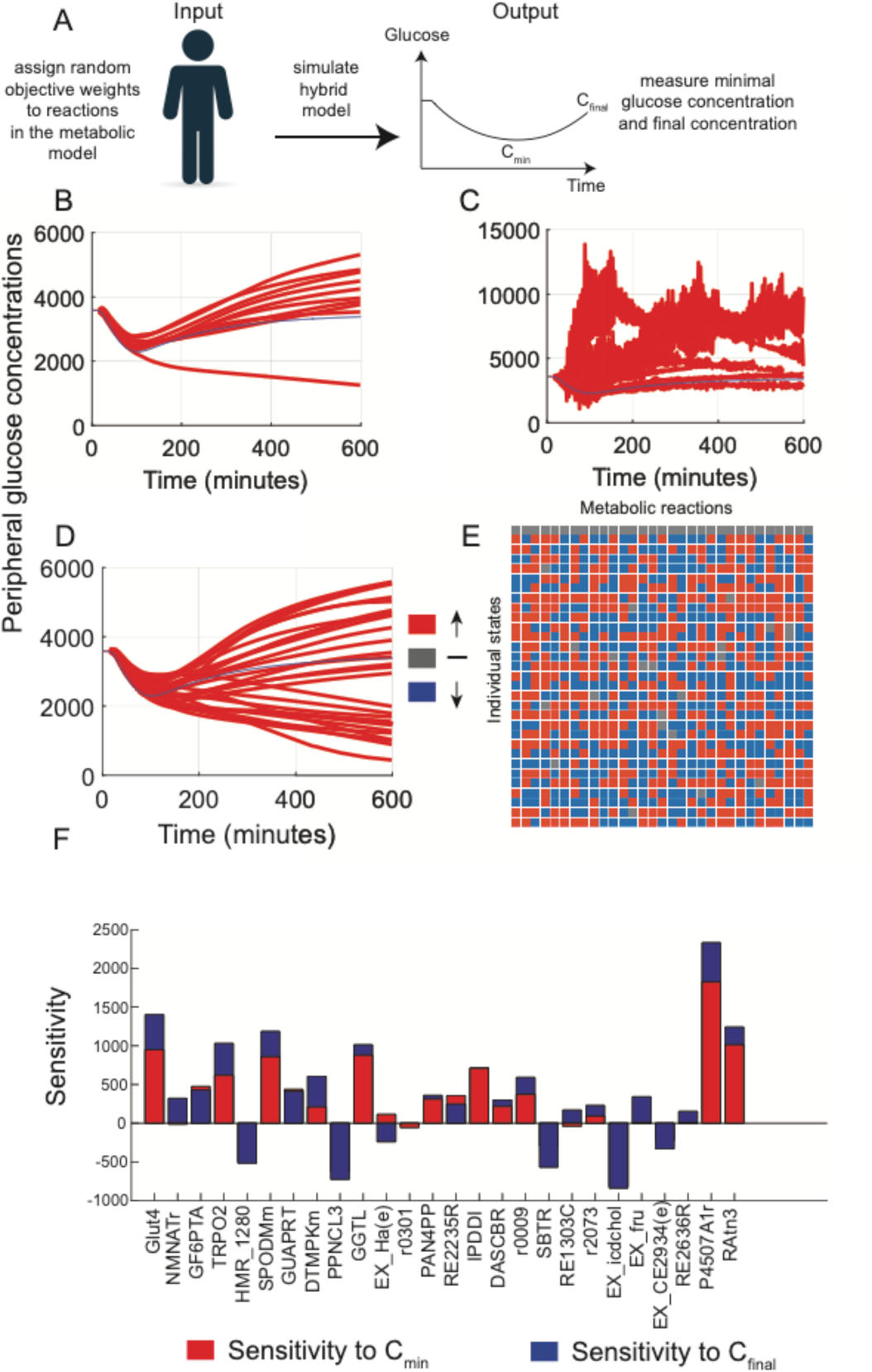
Intraindividual variability is assessed through a sensitivity analysis of the integrated model. **A**- The method consists of assigning random objective coefficients to metabolic reactions and measuring the minimum and final concentrations of glucose in each state with the kinetic parameters set to the population mean. **B**- Peripheral glucose profile when only the interface reactions are varied; the simulation is carried out with FBA (n=11). **C**- Glucose profile when both the interface reactions and metabolic model reactions are varied using FBA (n=11). **D**- Glucose profile when both the interface reactions and metabolic model reactions are varied using CRONICS (n=31). **E**- A 31-column excerpt of the matrix of the variation of the reaction weights in the objective for each state. The total number of considered reactions was 2,817. **F**- Computed sensitivities of each reaction in the internal state to the external output (minimum and final concentrations of peripheral glucose). The reactions are described in Table S7 and at http://vmh.life (Noronha et al., 2019).

The computed intraindividual variability was 30% *in silico* and was consistent with the empirical value of 12-45% (Heinemann, 2002). To determine the influence of each reaction on the metabolic profile of glucose, a multivariate regression was performed using the coefficient matrix as the input and the matrix of glucose concentrations per state as the output. The minimum and final glucose concentrations were considered in the sensitivity analysis as these concentrations inform about adverse reactions and treatment efficacy, i.e., hypoglycemia and hyperglycemia. The reaction sensitivities to both of the readouts (Figure 6-F) showed that GLUT4 transport contributed largely to the glucose profile. Interestingly, the reactions from the bile acid synthesis pathway also modulated the peripheral glucose profile. Taken together, these findings show that the internal reactions in the WBM model that were not necessarily in the interface reaction set could modulate the glucose concentrations in the GIM model through glycolytic and nonglycolytic pathways, thereby suggesting novel strategies to achieving diabetes control.

## Discussion

We developed a multiscale, dynamic, and organ-resolved model through the multialgorithm integration of metabolic and regulatory processes in T1D. The short timescale insulin-glucose-glucagon regulatory dynamics (GIM) served as constraints for the whole-body metabolic network (WBM). In addition to the expected target effects, the mechanism-unrelated metabolic effects of both insulin and T1D pathophysiology were included in the resulting diWBM model through organ-specific gene expression and metabolomics time-course data. The diWBM provided a complete picture of the network dynamics, regulation, and response to perturbations in relation to the known symptoms and clinically relevant variability in the insulin response, both within and between subjects.

### Chronic inflammation in type 1 diabetes improves disease identification using whole-body fluxes

To assess the impact of tolerance tests at the whole-body level, the WBM model was constrained by the glucose-insulin-glucagon regulation modeled by the GIM model in insulin and glucose challenges involving healthy and T1D states. The metabolic fluxes were subsequently used as features in the SVM binary classifier (Shaked et al., 2016). Identifying insulin versus glucose challenges at five minutes and 15 minutes after the perturbation was possible due to the opposing effects induced by these substances in the human body (Figure 2-C). The identification of the healthy and T1D flux distributions improved the separation between healthy and T1D individuals after the addition of gene expression constraints in T1D, representing the chronic inflammation that triggers the pathogenesis of the disease. While glycemia control is achieved through allopathy, the inflammatory aspect of the disease is often overlooked (McCall and Farhy, 2013) and must be equally addressed. Additionally, we could define the healthy and diabetic boundary using whole-body metabolic fluxes despite the subjected constraints affecting mainly the glucose-related pathways (Figure 2-D). These findings further reinforce the hypothesis that T1D is a multifactorial, pan-organ and systemic disease because of the key role that glucose plays in the regulation of metabolism and energy.

### Differentially regulated fluxes are suggestive of the mechanisms underlying the systemic symptomatology in type 1 diabetes

We used gene expression data from T1D pancreatic islets (Planas et al., 2010) as constraints to represent the pathophysiology of the disease in terms of both chronic inflammation and the disruption of glycolytic processes. The total number of metabolic reactions in T1D was the same as that in the healthy dWBM model, and there is no evidence for a complete metabolic gene knockout in affected patients. The dWBM model predicted a decrease in ATP in adipocytes (Figure 3-B) following the IVGTT test, as glycogen storage pathways are preferentially activated over ATP-producing pathways. The significantly decreased pathways in T1D mainly encompassed transport pathways in relation to the pan-organ distribution of insulin and the systemic properties of the disease. The well-known switch to fatty acid synthesis in T1D was demonstrated in the enrichment of corresponding pathways (Figure 3-E). In particular, the upregulation of sphingolipids has been linked to a decrease in tissue insulin sensitivity (Russo et al., 2013) in metabolic disorders and obesity. Interestingly, tyrosine metabolism was significantly ranked in the upregulated processes in T1D (Figure 3-E), and a recent study has identified direct links between diabetes and tyrosine pathway disruption (Ferguson et al., 2013). Additionally, an alteration in phosphoinositide metabolism has been reported in streptozotocin-induced diabetes in platelet cells (Jethmalani et al., 1994). Moreover, the glycosaminoglycan family (chondroitin, keratin sulfate, and N-glycan), which includes naturally occurring molecules that maintain tissue and cartilage, is decreased in T1D and could be fundamental to the long-lasting manifestations of this disease, such as the well-known disruption of tissue structure, e.g., diabetic foot (Yazdanpanah et al., 2015). A study in rats has shown a decrease in chondroitin sulfate in the kidney, suggesting possible implications of diabetes-induced nephropathy (Joladarashi et al., 2011). Collectively, the downregulation of tissue remodeling pathways added further evidence to the observed eschar-related symptomatology in diabetes. Since these manifestations occur during the later stages, the tissue remodeling pathway (Gowd et al., 2016) could be considered an interventional (either allopathic or nutritional) target during the diagnosis phase.

The enrichment of genes associated with the increased fluxes in T1D (Table S2) showed a representation of classical pathways, such as oxidative phosphorylation and carbon metabolism. Interestingly, Alzheimer’s disease (AD) and Parkinson’s disease (PD) were found to have potentially common links to T1D (Table S5, Figure S5). This finding further supports the growing evidence (Lalic et al., 2008; Moran et al., 2015) that neurodegenerative disorders and late-stage diabetes share a common pathogenesis (Cukierman et al., 2005). Furthermore, a clinical trial has repurposed Exenatide, a glucagon-like peptide-1 (GLP-1), in the treatment of PD (Athauda et al., 2017; Aviles-Olmos et al., 2013; Aviles-Olmos et al., 2014). In AD, a study has shown that higher plasma and brain glucose levels have been implicated in the disease progression (An et al., 2017) and the severity of the pathology possibly due to GLUT3 reduced brain glycolytic flux (Janson et al., 2004; Sims-Robinson et al., 2010). Similarly, antidiabetic drugs have been suggested for AD (Guney et al., 2016; Yarchoan and Arnold, 2014), particularly GLP-1 agonists and glucagon (Tai et al., 2018).

In addition, a drug repurposing analysis found that Mibefradil and Amlodipine could reverse the expression of the genes associated with the differentially active reactions in T1D (Table S6). Encouragingly, studies investigating experimental rodent models of diabetes have shown that calcium blockers improve blood glucose levels and diabetes-associated nephropathy (Lu et al., 2014; Ma et al., 2004). Taken together, these findings suggest the existence of a continuum between diabetes and neurodegenerative disorders, possibly involving a strong metabolic component.

### Insulin rewires interorgan exchanges

Conceptually, in the dWBM model, we added the dynamic features of the GIM model to the WBM model, enabling a hybrid modeling approach to metabolism. While the GIM model accurately predicted the glycolytic and regulatory effects of insulin, it could not capture the decrease in amino acids in the blood and other well-known anabolic effects (Dimitriadis et al., 2011). The addition of a dynamical model of insulin off-target effects (Yugi et al., 2014) to the diWBM model led to accurate predictions several known nonglycolytic insulin effects by the diWBM model (Figure 4). In particular, the prediction of the triglycerides time-course in the adipocytes (Figure S7) using diWBM showed an increase in as a result of the inhibition of lipases. Furthermore, we predicted the effects of insulin on metal homeostasis as metals play a prominent role in developing injection shocks due to the fast depletion of phosphate and potassium. Phosphate and potassium showed a trend towards an increase in uptake, although these results are not conclusive (Figure 4). Given the important role of these ions in signaling pathways and the demonstrated insulin-induced signaling modulation (Yugi et al., 2014), signaling pathways might play a greater role than metabolism in ion homeostasis. Considering the small concentrations of these ions compared to those of larger molecules, metabolism alone does not capture the full spectrum of micronutrient homeostasis.

Finally, to study the effect of insulin on interorgan crosstalk, a subset of reactions was selected to encompass the kidneys, liver, brain, pancreas, muscle, adipocyte, and interorgan compartments, e.g., plasma, as the role of these organs has been demonstrated in the disease pathology (Li et al., 2009; Romacho et al., 2014). The total interorgan crosstalk increased as an effect of the insulin administration (Figure S8), further structuring the organs in a metabolic continuum. The multiple organ complications related to late-stage diabetes corroborate this finding as the loss of insulin mediates the decrease in organ crosstalk at a metabolic level. Although the implications of endocrine secretions have been more thoroughly studied in the pathology of systemic organ failure (Li et al., 2009), the overall decrease in organ crosstalk in T1D might be a combined effect of the signaling and metabolic properties of insulin.

### Insulin-mediated hysteresis reflects between-subject variability

The generation of a synthetic population of T1D patients involved the variation of patient-specific kinetic parameters (Schaller et al., 2013). The obtained peripheral glucose profile reproduced the reported 25-35% interindividual variability in the insulin response, and the differences were particularly notable in the AUC, C_min_, and T_min_ (Figure 5-A, B). Interestingly, the patients had a similar number of active reactions, reflecting a similar network topology (Figure 5-C), which was an effect of the simulation framework CRONICS that ensured a sparse set of fluxes and a minimal change between time intervals. This observation of similar network topology also implies that the constitutive metabolic background is similar among the patients and that the high-flux backbone is conserved (Almaas et al., 2004) in the absence of a specific enzyme deficiency as in the case of IEMs (Sahoo et al., 2012). However, by projecting the principal components of the flux distribution of individual patients during the simulation time, a metabolic shift could be observed (Figure 5-D). Thus, two main results were deducted. First, the differences in the response to insulin was mainly due to changes in flux values in the metabolic model (Figure 5-D) rather than the network structure. Second, the final metabolic state differed from the initial state, although the simulation time was sufficiently long (10 hours) to ensure a return to the steady state in the GIM model uncoupled from the WBM model, which is consistent with the profound insulin-induced regulatory mechanisms in metabolism. Each patient achieved a differential control of the metabolic shift, which could be related to the varying metabolic outcomes (Figure 5-E). In particular, insulin, which was indirectly represented by the peripheral glucose levels, induced a hysteresis in metabolism (Figure 5-F), which highlights the dramatic metabolic changes that occur following insulin administration. In fact, the binding of insulin to its receptor and the release of GLUT transporters with an ∼15 hour half-life under insulin treatment (Sargeant and Paquet, 1993) can be key drivers of the shift in the metabolic steady state given the ubiquitous distribution of insulin receptors. Moreover, the hysteresis reflected the correspondence of several internal metabolic states to a unique glucose concentration (Figure 5-F). This finding corroborates the unreliability of plasma glucose levels as a universal marker for the diagnosis of diabetes (Bonora and Tuomilehto, 2011). Additionally, the observed hysteresis revealed a differential action of insulin in the population, and for most patients, the hysteresis described a quasicycle of glucose concentrations predicted by the GIM model and the internal system properties modeled by whole-body metabolic fluxes. Notably, a few virtual patients did not achieve a metabolic shift and remained at the initial state (Figure 5-F), denoting ineffective insulin action, while a large fluctuation into the final state can be a marker of hypoglycemia or hyperglycemia and uncontrolled diabetes.

Finally, as we demonstrated earlier, the values of the metabolic fluxes, rather than the network structure, were the drivers of the observed differences among the synthetic patients (Figure 5-G). In particular, the citric acid cycle and oxidative phosphorylation supported the observed between-subject variability, while glycolysis and the pentose phosphate pathways were resilient to the perturbation, highlighting their essential role in human metabolism. In the architecture of metabolism, glycolysis and the pentose phosphate pathway occupy a central position, while the citric acid cycle and oxidative phosphorylation are the entry points to several secondary redundant pathways. In fact, external network nodes tend to act as buffers that absorb the perturbations and maintain the central functionalities of the network (Gilarranz et al., 2017). Accordingly, a reduction in interindividual variability in response to insulin is key to achieving diabetes control. Notably, a study involving a cohort of 20,303 individuals (Akirov et al., 2017) has shown that a higher coefficient of variance of glucose dynamics in diabetic patients has been correlated to higher mortality rates; therefore, reducing the within-patient variability is an additional major determinant of the management of T1D.

### GLUT4 as a pharmacogenomic target for diabetes control

The within-patient variability in the insulin response poses a great challenge for the identification of the internal factors that could modulate the dynamics of glucose. Assuming that the same kinetic parameters represented the average patient, we randomly generated several internal metabolic states and subsequently, computed glucose time series after insulin administration. Each state assumed different objective weights of a selected set of reactions. Moreover, CRONICS ensured the smoothness of the simulated system, and in contrast to the simulation setting of the interindividual variability, the outcome of the constraint-based model determined the dynamics of glucose. The outcome of the simulation was consistent with the empirical intraindividual variability in the insulin response (Heinemann, 2002), and the reactions were subsequently classified by their sensitivity to the final and minimum glucose concentrations, which acted as surrogate endpoints of insulin activity (Figure 6-F). Given the high computational cost of the simulations, we randomly selected representative reactions from each subsystem in all organs. Although the number of core interface reactions shared between WBM and GIM was limited, the modulation of reaction weights in WBM could influence metabolite kinetics in GIM through trans effects but with variable sensitivity. The bile acid synthesis pathway was found to highly modulate glucose concentrations in the model. Studies have shown that dysregulation of the pathway could contribute to the pathogenesis of diabetes through the modulation of GLP-1 and insulin-sensitizing activities (Prawitt et al., 2011; Tomkin and Owens, 2016). Additionally, the transporter GLUT4 had a high sensitivity towards the minimum and final glucose concentrations. The insulin-dependent carrier was released to balance high glucose concentrations, and its expression could be a major factor in the pharmacodynamics of insulin (Correa-Giannella and Machado, 2013). Generally, the transport subsystem, which is mediated by carrier proteins, provides a rationale for the design of novel antidiabetic drugs targeting transport proteins, such as SGLT1 (Sands et al., 2015) and SGLT2 (Verma et al., 2017). Finally, since the flux through the GLUT4 reactions highly modulated the glucose concentrations in the diWBM model, we hypothesize that the carrier could be a major pharmacogenomic effector of insulin action. The design of adjuvant therapy of insulin targeting the expression and release of GLUT4 is a potential avenue for decreasing the intraindividual variability in the insulin response, overcoming insulin resistance, and ultimately achieving diabetes control. Overall, the diWBM model presented in this study expands our understanding of T1D and has the potential to empower evidence-based approaches to human pathology.

## Acknowledgments

The authors would like to acknowledge Bayer Technology Services for providing an academic version of the PK-SIM/MOBI software suite and the source file for the GIM model, Mr. Katsuyuki Yugi from the University of Tokyo for providing the insulin model, and the Molecular Systems Physiology lab members at the University of Luxembourg for reviewing the manuscript and valuable discussions.

## Author contributions

MBG and IT conceived the study. MBG developed the framework and carried out the simulations and analysis. MBG and IT wrote and edited the manuscript.

## Supplemental information

The supplemental information includes nine figures and seven tables.

## Data availability

All related code will be available in https://github.com/ThieleLab upon publication

## Methods

### Coupling the dynamical model and constraint-based model

The male WBM model, version 1.00 (May 2017), is an organ-resolved metabolic model of the human body that encompasses 22 organs and six blood cells totaling more than 80,000 reactions and 50,000 metabolites (Thiele et al., 2018). The organ specific reactions were derived through the curation of evidence from the literature and high-throughput experiments, such as proteomics and metabolomics data (Thiele et al., 2018).

The metabolic reaction fluxes in the WBM model were calculated using constraint-based modeling (Orth et al., 2010) by solving the following linear program:

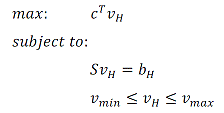

where *c* is the objective coefficient vector; *S_m,n_* is the stoichiometric matrix of *m* metabolites and *n* reactions; and *v_H_* is a flux vector bounded by *v_min_*, which is the lower-bound vector, and *v_max_*, which is the upper-bound vector. When *b_H_* = *o_m_*, the problem is constrained by *S_v_* = 0, also referred to as FBA (Orth et al., 2010). The bounds of the reaction rates in the WBM model were derived from physiological parameters, such as blood flow and clearance rates, in addition to thermodynamics and metabolite concentrations (Thiele et al., 2018).

The GIM model is a physiology-based pharmacokinetic (PBPK) model of the regulation of glucose by insulin, glucagon, and incretins. The GIM model covers 20 organs and models the complex dynamical processes of insulin receptor binding and internalization, the subsequent release of glucose transporters, and the detailed subcompartments, including the interstitial space and endothelium (Schaller et al., 2013). In the intestinal tract, the GIM model models the intestinal regulation of glucose by incretin hormones (glucagon-like peptide 1 (GLP1) and gastric inhibitory polypeptide (GIP)) and the absorption by glucose carriers, such as SGLT1 and SGLT2. In each organ, particularly the target organs, i.e., liver, pancreas, brain, fat, and muscle, the GIM model represented a number of processes related to the action of insulin, such as its diffusion from the blood into the extracellular space, the metabolism of glucose by red blood cells, the internalization of insulin by endothelial cells, the clearance by the lymphatic circulation, the binding and internalization of insulin receptors followed by the release of glucose receptors, and the storage of glycogen. Since insulin acts at a relatively low dose, its binding the receptor can modulate its pharmacokinetics. Consequently, the GIM model modeled the target-mediated drug disposition (TMDD) of insulin in relation to its pharmacodynamics. The organs modeled included bone, brain, fat, gallbladder, gonads, heart, kidney, large and small intestine, liver, lung, muscle, pancreas, skin, spleen, and stomach connected by extracellular compartments and biofluids, such as arterial and venous blood, portal vein, intestinal lumen, and saliva. Each organ is further compartmentalized into anatomical and histological units, such as the intracellular space, the interstitial space, and the endothelial compartment. The overlapping organs between the WBM model and the GIM model are the lungs, brain, heart, stomach, colon, small intestine, skin, testis, spleen, kidneys, adipocytes, muscle, liver, pancreas, and blood cells.

The points of intersection between the dynamical model (GIM) and the constraint-based model (WBM) included metabolites and reactions. There were four cases (Figure 1) resulting in different implementations of the coupling constraints (see also Methods-Construction of the hybrid model). Let us first consider the concentrations of a given metabolite modeled in the GIM model (G) by the following ODE:

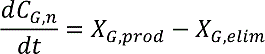

where *X_prod_* and *X_elim_* are reaction fluxes producing and eliminating metabolite *n,* respectively, through first-order processes.

*Case 1:* If one reaction in the dynamical model corresponded to one reaction in the metabolic model, the following constraints were implemented for one time step:

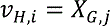

where *t* is time; *v* is the flux vector; *i* is the index of the reaction in the WBM model (*H*); and *j* is the index of the reaction in the GIM model, such that reactions *i* and *j* perform the same metabolic function, e.g., the liver hexokinase reaction.

*Case 2:* When a metabolite was presented in both models, the sum of its anabolic and catabolic fluxes in the WBM model was constrained by its change-of-concentration in the GIM model, which is also referred to as metabolite pooling fluxes (Covert et al., 2008). The constraints were formulated as follows:

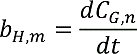

where *m* refers to the metabolite in the WBM model, *n* refers to the metabolite in the GIM model, *C_n_* is the concentration of *n*, and *b* is the vector of metabolite change-of-concentration in the WBM model.

*Case 3:* One reaction in the GIM model was represented by more than one reaction in the WBM model. Typically, these reactions were catalyzed by cofactor-dependent enzymes, such as hexokinase, with ATP and ADP as cofactors. The reaction pooling fluxes were treated as follows:

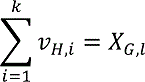

where *v_1_* to *v_k_* are the reactions in the WBM model and correspond to reaction *l* in the GIM model.

*Case 4:* One metabolite in the GIM model corresponds to more than one metabolite in the WBM model. For example, red blood cell glucose in the GIM model corresponds to glucose in red blood cells, monocytes, natural killer cells, B cells, platelets, and CD4 T cells in the WBM model. The constraints were formulated as follows:

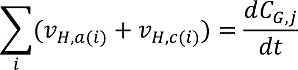

where *v_a_* refers to the anabolic fluxes, *v_c_* refers to the catabolic fluxes, and *i* is the index of different metabolites in the WBM model corresponding to metabolite *j* in the GIM model.

After implementing these constraints, the steady state was assumed in the chosen time step for the remaining, nonoverlapping metabolites, such as amino acids, present in both models. Consequently, the constraints translated as follows:

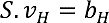

where *S* is the stoichiometric matrix in the WBM model, *v* is the flux vector, and *b* is the metabolite concentration change that equaled zero for the steady-state metabolites and nonzero for metabolites belonging to case 2. When the applied constraints rendered the linear program infeasible, the lower and upper bounds were minimally relaxed in both the amplitude of relaxation and the cardinal of the reactions to be relaxed (Methods-Relaxing infeasible problems) (Fleming and al., Submitted).

The simulation of the tolerance tests and the assessment of the inter and intraindividual variability in the insulin response were performed using the dWBM model with specific model parameters. The tolerance test parameters were the same as those reported for the GIM model (Schaller et al., 2013) (Table S3), which included trials of intravenous insulin tolerance test (IVITT), intravenous glucose tolerance test (IVGTT), baseline glucose concentration (Noinf), subcutaneous insulin bolus (SCIB), subcutaneous insulin infusion (SCII), solid meal (WB-Solid), and oral liquid glucose solution (WB-Liquid) in healthy and T1D patients. The parameters of the GIM model were reportedly (Schaller et al., 2013) identified using tolerance test trial data (Regittnig et al., 1999; Sorensen, 1985) and bihormonal closed-loop experiments (El-Khatib et al., 2010). The WBM model was simulated using exchange reactions corresponding to a standard European diet (Thiele et al., 2018).

### Construction of the hybrid model

Coupling the GIM model with the WBM model fell under one of the following scenarios (Figure 1):

1. When the fluxes in the GIM model and the WBM model corresponded exactly, the flux in the metabolic model was set to that in the dynamical model.
2. When the metabolites were identical in both models, the corresponding *b* vector entry in the metabolic model was set equal to the rate of change in metabolite concentration.
3. When a specific reaction in the dynamical model corresponded to several reactions in the metabolic model, the reactions were pooled in the WBM model, such that their total flux equaled the reaction flux in the dynamical model. For example, this case applies to the hexokinase isoforms.
4. Finally, when a specific metabolite in the dynamical model corresponded to several metabolites, the metabolites were pooled into one supermetabolite by merging the corresponding rows in the stoichiometric matrix. In the dynamical model, the blood cells were pooled into one compartment, but in the metabolic model, each blood cell was represented separately.

### Simulation setting

#### Tolerance tests

We used the dWBM model to simulate the outcome of different tolerance tests. The kinetic parameters provided with the GIM model were used to represent the doses and time of intake. We dynamically applied the constraints in the WBM model in each time step following the indirect coupling method. In IVGTT, a dose of glucose is injected intravenously, hence, we added an exchange reaction to represent the intake of glucose in the blood *glc_D[bc]* during the 15 minutes of infusion (67.7 mmol/5 mn). Subsequently, we solved the WBM model using parsimonious FBA (pFBA) (Lewis et al., 2010) assuming a whole-body maintenance objective function and aggregated the results in a matrix for each time step comprising the tolerance tests in the columns and the metabolic fluxes in the rows. We reduced the matrix through PCA and plotted the time course of the first component (Figure 2-E) to illustrate the whole-body metabolic shift induced by insulin and the glucose tolerance tests. The points were fitted to a 6^th^ order polynomial curve (Figure 2-E).

Furthermore, to assess whether the dWBM model was sensitive to the dynamical constraints applied from the GIM model, we aggregated the pFBA simulation results over 600 minutes of simulation per five minute time steps. Using a PCA, we reduced the dimensionality of the predicted fluxes to three dimensions, which captured 95% of the variability with the dWBM models representing only the glucose-insulin-glucagon effects (target effects) (Figure 2-C). The models additionally including differentially expressed genes (off-target effects) captured 93% of the variability in T1D (Figure 2-D). In each tolerance test simulation, we built a feature matrix comprising the whole-body metabolic fluxes in the columns and the tolerance test in each time step in the rows. At a time step of five minutes, 120 simulation results in the T1D and healthy states formed the 240 rows. Then, we reduced each matrix of the tolerance test using PCA. We considered the first ten PCA components that had *p<0.001*. The component significance was assessed through 100 random permutations of the columns, followed by the PCA of the perturbed matrix. The first ten components were then used as features in a binary SVM to classify the healthy and T1D states based on the whole-body fluxes in each tolerance test. We used an SVM with a Gaussian kernel, performed 3-fold cross-validation on the training set (80%) and predicted the labels of the test set (20%). The data were standardized, and the process was repeated 100 times considering each time a different partition of the training and test set.

We chose to use the significant components as features instead of performing feature selection on the whole-body fluxes as the main question was the assessment of the dWBM model sensitivity towards glucose and insulin challenges and the subsequent whole-body metabolic shift. Furthermore, the activated reaction fluxes in each time step are among many possible solutions in the AOS space and cannot be used as conclusive evidence of the disrupted metabolic pathways in T1D and healthy models, even though pFBA considerably reduces the AOS space (Toroghi et al., 2016). We addressed the question of disrupted metabolic pathways in T1D by performing FVA in T1D and healthy models and using all solution vectors of the AOS as a basis for comparison. The dWBM model was predictive of the T1D and healthy states in the tolerance tests (Figure 2-E); moreover, when we used insulin and glucose challenges as classes in the SVM instead of the T1D and healthy states, the whole-body fluxes were also predictive of the states (Figure S2). Taken together, the dWBM model is predictive of both the condition (T1D and healthy state) and the perturbation (insulin and glucose challenges).

### Relaxing infeasible problems

Subjecting dynamical model-derived constraints to the metabolic model could result in unfeasible problems. This situation could arise from conceptually different modeling approaches. In fact, the GIM model depicted short time events, while the constraint-based model was designed to simulate steady states. For instance, while organs, such as the lungs, are commonly known to be glucose metabolizers rather than consumers, the GIM model could show a small secretion of glucose in a short time step as a consequence of free diffusion through the organ membrane. We relaxed the corresponding reactions in the WBM model to obtain a feasible model by minimally relaxing the upper and lower bounds of the internal reactions. If the standard linear program is infeasible,

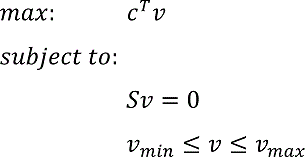

where *c^T^v* is the objective function, *v* is the flux vector of the metabolic reactions, *c* is the vector of the objective coefficients, *S_(m,n)_* is the stoichiometric matrix linking *m* metabolites and *n* reactions, *lb* is the vector of the reaction lower bound, and *ub* is the vector of the reaction upper bound. The following problem minimally relaxes the infeasible model (Fleming and al., Submitted):

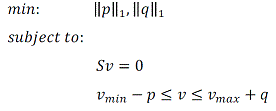

where *p* is the relaxation vector of the lower bound, and *q* is the relaxation vector of the upper bound. Minimizing the 1-norm of *p* and *q* ensures sparsity (a minimal cardinal of reactions to be relaxed) with a minimal total sum of relaxation amplitude.

### Comparison of the predictive capabilities of the different models

To compare the predictive capabilities of the WBM, the GIM, and the dWBM models, we simulated an IVGTT. The GIM model includes a parameter set for simulating IVGTT (Table S3), and the glucose concentrations were obtained by integrating the ODEs. The glucose concentrations in the WBM model were obtained through dynamic FBA (dFBA). Briefly, the initial amount of glucose was set as availability constraints and decreased in each time step after subtracting the consumed glucose fluxes obtained through solving the linear program (Mahadevan et al., 2002). The maximum rate of change of glucose was set to the intravenously injected amount in the *b* vector; then, the problem translates as follows:

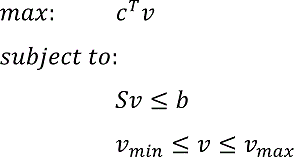

The dWBM model not only reproduced the glucose time series of the GIM model but also could predict the time course of ATP, which was not present in the GIM model. The simulation was performed by setting the ATP demand reaction (VMH ID: DM_atp_c_ (Noronha et al., 2019)) at each time step as the objective function as follows:

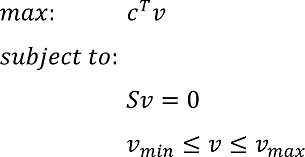

where *c* is the vector of objective function coefficients; and *v_min_* and *v_max_* are the minimum and maximum fluxes through a reaction, respectively. The *c* vector entry corresponding to the indices of reactions corresponding to the ATP demand in each organ was set to one. Then, the baseline value of the ATP demand reaction flux, which was calculated at the initial simulation time, was subtracted from the obtained value. The cumulative sum of the resulting fluxes enabled the theoretical time course of ATP in a given tissue to be obtained.

### Modeling T1D off-target effects in the pancreas

We considered T1D as a combination of target and off-target effects. The target effects are directly related to the decrease in insulin secretion and the increase in glucose levels, and the off-target effects are represented by the underlying chronic inflammation that triggers the of disease. To model the target effects, we constructed healthy and T1D dWBM models that were obtained by coupling the WBM model with the healthy and T1D GIM models. Moreover, gene expression data (Planas et al., 2010) from pancreas biopsies T1D patients obtained through whole transcriptome sequencing of four patients after five days, nine months, five years, and ten years postdiagnosis were used to further constrain the dWBM diabetic model and capture chronic inflammation leading to the development of T1D in the pancreas. The differentially expressed genes were mapped to the metabolic model to represent the off-target effects of the disease. Among the list of 475 differentially expressed genes, only the metabolic genes present in the WBM model, corresponding to 24 genes, were kept for further analysis (Figure S3, Table S1). Consequently, the bounds in the linear program of the dWBM T1D model were modified in the reactions corresponding to each gene such that

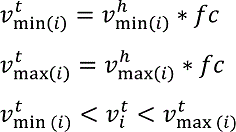

where *fc* is the gene expression fold change between the control and disease of the gene encoding reaction *i*, *v^h^_min(i)_* is the minimum reaction flux corresponding to reaction *i* in the healthy model, and *v^h^_max(i)_* is the maximum reaction flux in the healthy model as identified by the flux variability analysis. *v^h^_min(i)_* and *v^h^_max(i)_* are the new lower and upper flux bounds computed in dWBM type 1 diabetic models. The gene expression data constrained a total of 80 reactions in the pancreas (Table S2).

### Modeling off-target insulin effects in the liver

Similar to T1D, we considered insulin to induce target and off-target effects. The target effects of insulin include the reduction in glucose levels, and the off-target effects include various physiological processes involving insulin, such as the regulation of glycolysis. The off-target effects of insulin in the liver were modeled by coupling an additional ODE model to the dWBM model, resulting into the diWBM model. The additional ODE model (Yugi et al., 2014) represented the metabolic effects of insulin on the glycolytic fluxes in one hour. The amounts of eight metabolites *in vitro* (fructose-6-phosphate, fructose-1,6-diphosphate, phosphoenolpyruvate, isocitrate, 2-oxoglutarate, malate, fructose-2,6-biphosphate, and citrate) were scaled to the amounts (mmol) *in vivo* as follows:

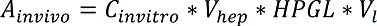

where *V_hep_* is the hepatocyte volume, *HPGL* is the hepatocellularity, *C_invitro_* is the concentration of the metabolites *in vitro* and *V_l_* is the liver volume.

The values of the parameters were set as follows (Heinemann et al., 1999):

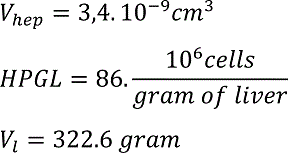

The constraints were applied by setting the metabolites’ rates of change in the insulin dynamical model equal to the corresponding *b* vector value in the WBM model.

### Simulation of interindividual variability in the insulin response

To assess the between-subject variability in the glucose dynamics after a bolus of insulin (Table S3), the identified differential parameters (Schaller et al., 2013) (Table 1) between the T1D and healthy GIM models were randomly varied within a 2-fold interval to reproduce the observed 25-35% variability in the patient population (Heinemann, 2002) as computed by 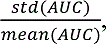 where *AUC* refers to the area under the curve of the peripheral glucose concentrations per patient. The parameters were randomly increased or decreased for a group of 30 virtual patients, and the coupling between the newly obtained GIM model and the WBM model was performed as previously described.

### The simulation algorithm CRONICS

The simulations were performed according to the following two main methods: direct and indirect coupling (Covert et al., 2008; Krauss et al., 2012).

*Indirect coupling* was used to dynamically constrain the WBM model with respect to each time step. First, the GIM model was simulated for the entire time horizon, and the reaction rates were computed and applied retrospectively as constraints in the WBM model as described above (see Methods-Coupling the dynamical model and constraint-based model, Figure 1). In this case, the WBM model represents the pharmacodynamic endpoint of the GIM model.

*Direct coupling* assumes an interdependency of both models, where the fluxes of the GIM model for a specific time step were applied as constraints in the WBM model, and the result of the linear program defined the initial rates in the GIM model for the consecutive time step. Here, the outcome of the simulation in the GIM model was dependent on WBM and vice-versa. In this case, the WBM model influenced the pharmacokinetics of the GIM model.

The integrative framework CRONICS (Figure S4) included the coupling scenarios mentioned below (Methods-CRONICS framework) and additionally ensured the smoothness of the hybrid model. In fact, given the high dimensionality of the WBM model, there can be more than one solution to the optimal objective of the linear program, which forms the alternate optimal solution (AOS) space. Therefore, in each time step, the solution of the linear program can be distant from the previous solution, which compromises the smoothness of dynamical simulations in large-scale metabolic models. We addressed this issue by i) performing pFBA in each time step to obtain a reduced AOS space as previously suggested (Toroghi et al., 2016); ii) minimizing the Euclidean distance between the flux vectors in each time step similar to using Minimization Of Metabolic Adjustment (MOMA) (Segre et al., 2002); and iii) optimizing a set of reactions in each time step as previously suggested (Gomez et al., 2014). The latter was used to simulate the intraindividual variability in the insulin response. Additionally, the solver parameters had to be tuned to optimize convergence and ensure feasibility (Methods-Software and solver parameters).

### CRONICS framework

The simulation of the hybrid model faced the following challenges:

- Given the size of the solution space in each time step and the effect of alternate optimal solutions, the problem is largely underdetermined.
- The large size of the metabolic model required a reduction in the set of active reactions in the subsequent analysis and biological interpretation.
- The simulation time, as a model could typically run for 2-3 days for simulations representing a 10-hour time window.

The abovementioned challenges were addressed by combining a mosaic of existing techniques in the CRONICS framework, which consisted of the following steps:

1. (optional) Depending on the simulation, the solution basis of the unconstrained problem was generated before coupling the dynamical and metabolic models. Then, the solution basis was set in all subsequent optimization problems, while activating the advanced start option in CPLEX as follows:

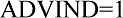
2. The setting started with the simulation of the dynamical model for one time step. Based on dFBA, the computed constraints were subjected to the metabolic model, and a 1-norm minimal (pFBA) solution was selected given its sparse properties, which allowed for the selection of a smaller set of active reactions for further analysis.
3. (Indirect coupling) The following time step in the dynamical model was simulated, and the constraints were computed and subjected to the metabolic model. The selected solution was the solution that was minimally distant to the previous solution (MOMA). This process guaranteed the smoothness of the hybrid system and the chronology of the simulation. Although the simulation was computationally expensive due to the large size of the solution vector, we chose a subset of the vector to minimize. The minimal subset could be the coupled reactions; in this case, the obtained vector was indeed nearly identical to the initial subset vector, which resulted in all simulations being converged to the dynamical model behavior in terms of smoothness, while the uncoupled reactions could abruptly change values in time steps as small as five minutes. A subset containing reactions of the brain, liver, kidney, adipocytes, muscle, and pancreas was allowed to globally constrain the fluxes with respect to the chronology of the simulation while maintaining the smoothness of the system.
4. (Direct coupling) In this previous setting, the hybrid model consisted of a dynamically constrained metabolic network. In the direct setting, the dynamical model was, in turn, constrained by the metabolic model. The obtained fluxes from the metabolic model served as inputs for the next time step of the dynamical model. If the solution fluxes corresponded to the input constraints, the hybrid model reached the behavior of the dynamical model alone. In both settings, the solution of the T1D model for the first time step was minimally distant to the healthy model for the first time step.

Due to the long simulation times, the simulation of the dWBM model using the CRONICS framework restricted the distance minimization of the flux vectors between the different time steps (Figure S4-step VI) to a subvector, including the pancreas, kidney, adipocytes, muscle, liver, brain, and fluids, e.g., peripheral blood, portal vein, blood-brain barrier, and intestinal compartments.

We used the CRONICS framework to predict the metabolite concentrations in the dWBM model. The metabolites fell into the following two categories: the metabolites whose time course was predicted by the GIM model and the metabolites in the WBM model that obeyed the steady-state assumption. Naturally, we cannot predict the concentrations of the metabolites that were in steady-state. Nevertheless, to predict the dynamic organ requirements of those metabolites, we created organ-specific demand reactions, which allow accumulation and depletion of metabolites thereby allowing to compute concentration change over time. We applied this method to predict the time course of ATP demand in the liver (Figure 3-B), the triglyceride demand in the adipocytes (Figure S7), and the valine and phenylalanine demand in plasma (Figure S6).

### Metabolic network topology

To analyze the topological features under different sets of perturbations, the WBM model, which is a hypergraph of metabolism, was transformed into a metabolite-centric graph, and redundant metabolites, i.e., *ATP*, *ADP*, *H_2_0*, *NH_4_*, *H^+^*, *NADH*, *NADPH*, and *Pi,* were removed to facilitate the subsequent analysis. Metabolic graphs have been previously shown to be scale-free networks endowed with a modular organization (Gomez et al., 2014). The distribution of metabolite connectivity across the metabolic network was accordingly fit on a power law as follows:

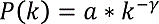

where *k* represents the metabolite degree, *a* and γ are the power law parameters, and *P(k)* is the relative frequency of each metabolite.

### Sensitivity analysis of the hybrid model, intraindividual variability, and multivariate regression

To assess the intraindividual variability in the response to the insulin injection, the GIM model parameters were fixed based on the mean patient anthropomorphic parameters (Schaller et al., 2013), and the flux values of a set of 2,817 randomly selected reactions representing all subsystems in the organs in the WBM model were allowed to vary by assigning random objective coefficients. Subsequently, a matrix, *X_(p,q)_*, of objective coefficients for each reaction was randomly generated, where *p* is the number of trials representing the within-patient metabolic states, and *q* is the number of reactions.

The output of the simulation of each trial with the corresponding objective coefficients was the concentration of glucose after the subcutaneous insulin injection at each of the *n* time steps. The vector *Y_(p,n)_* was the obtained as output vector and contained the concentration values at each time step. Using multivariate regression, as suggested previously (Sobie, 2009), the sensitivity vector ρ, representing the contribution of each reaction flux in the WBM model to the time course of the metabolites in the GIM model, was computed as follows:

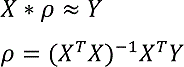

Then, the ρ vector allows for the quantification of the sensitivity of the peripheral glucose concentration to the considered metabolic reactions. Particularly, the minimum concentration *C_min_* and final concentration reached at the end of the simulation *C_end_* were considered for further analysis.

### Intraindividual variability in the insulin response

We modeled the interindividual variability in the insulin response as the variation in the kinetic parameters in the GIM model. Therefore, we created 31 GIM models and coupled them to the WBM model. The glucose concentrations were completely determined by the GIM model, which was also referred to as indirect coupling (Krauss et al., 2012). The modeling of the intraindividual variability in the insulin response consisted of using the average patient kinetic parameters in the GIM model and the variation in the internal state of the system as represented by the WBM model. In this case, we randomly selected 1% of the reactions in each subsystem in each organ, which resulted in a set of 2,817 reactions that equally represented all metabolic subsystems in all organs. We assigned each reaction a random objective weight, which corresponded to a metabolic state. Subsequently, we created 31 metabolic states and simulated the models after a subcutaneous injection of insulin. The glucose concentrations depend on both the WBM and the GIM model, which is also referred to as direct coupling (Krauss et al., 2012). The input matrix *X_p,q_* represents the objective coefficients of each *q* reaction in the *p* metabolic state. Here, *p*=31 and *q*=2,817. The output matrix *Y_p,n_* represents the glucose concentrations at each of the *n* time steps in the *p* metabolic states, with *n*=234 at a 2.5 minute time step length, and the infusion started at 16 minutes for a 10-hour simulation. Using a multivariate regression, we estimated the matrix *ρ_q,n_*, which represents the sensitivities of each *q* reaction to each *n* time step in the simulation. In particular, the time steps corresponding to the minimum concentration *C_min_* and the final concentration *C_final_* were considered for further analysis as these concentrations could be linked to hyperglycemia and hypoglycemia and diabetes control in general. The estimation of the sensitivity matrix consisted of solving the following equation:

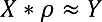

We used the *mvregress* routine in MATLAB and the covariance weighted least square (CWLS) algorithm to estimate ρ. The algorithm provides *p* coefficients corresponding to *p* of *q* reactions, and the remaining reactions are set to zero.

### Software and solver parameters

The simulations were carried out in MATLAB (2014b, Natick, MA, USA) with the ODE15s built-in function, COBRA toolbox v3.0 (Heirendt et al., 2019) and the CPLEX and MAD TOMLAB v7.9 implementation on Windows 7 professional and Ubuntu 16.04-based high-performance computing units. The following parameters allowed a faster convergence and resolving ‘infeasible after unscaling’ types of issues:

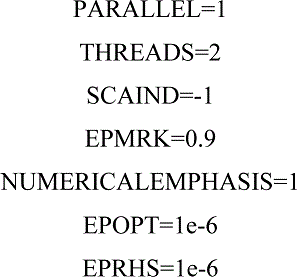

### Differential reaction fluxes between healthy and type 1 diabetic models

To compare the fluxes between the healthy and type 1 diabetic models, the comparison of individual fluxes could yield inaccurate results given the large alternative optimal solution space. We compared the flux distributions per reaction rather than the value provided by one solution. To obtain the flux probability distributions, we performed a flux variability analysis of both the healthy and type 1 diabetic model. The COBRA toolbox function *fastFVA* (Gudmundsson and Thiele, 2010) was used with the solutions as output as follows:

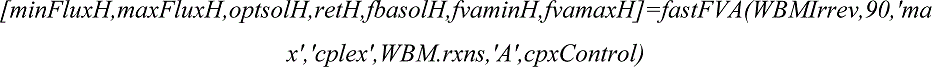

Prior to the flux variability analysis, the metabolic model was translated to its irreversible version such that each reversible reaction is decomposed in the forward and backward reaction in a way that the obtained fluxes are positive. The conversion was performed using the following COBRA toolbox function:

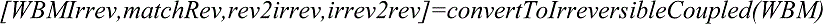

The two matrices containing the flux values per irreversible reaction in the healthy and T1D models were used as the input in the volcano plot function *mavolcanoplot* in MATLAB (2014b, Natick, MA, USA). A p-value less than 0.001 was considered the significance threshold for a minimal fold change of 1.3. Higher fold change values are usually used in gene expression experiments with the objective of eliminating small-value changes that are related to the batch effects and the experimental set up as well as other factors that introduce stochasticity. Conversely, using the same solver settings, linear programming solutions are deterministic, hence, we considered a smaller fold change value. After performing differential flux analysis, the irreversible reactions were mapped to their original identifiers to determine the impacted reactions, their subsystems, and their encoding genes.

### Enrichment of gene vectors

After obtaining the upregulated and downregulated fluxes in T1D models, we connected the reactions to their encoding genes using the defined gene-protein-reactions associations (Thiele and Palsson, 2010) and performed a gene set enrichment analysis. As the set of obtained genes could experimentally correspond to differentially expressed genes, we queried i) the LINCS database (Duan et al., 2014) to search for small molecules that reverse the signature of T1D, i.e., compounds that induce a genetic signature opposite to the T1D signature we obtained, which can potentially reverse the metabolic profile and ii) the KEGG database (Kanehisa et al., 2017) to assess which diseases were similar in signature to T1D.

To identify potential small molecules that could reverse T1D, we collected a list of upregulated and downregulated genes in T1D and queried the LINCS Canvas Browser (Duan et al., 2014), setting the upregulated genes and downregulated genes in the up and down fields, respectively, and using the *reverse* option to search for small molecules that reverse the queried signature. The results are reported in Table S6.

To identify diseases with a similar genetic signature to T1D, we merged the upregulated and downregulated reactions into a unique set of differentially active reactions. Since one metabolic reaction can be encoded by one or more genes using Boolean operations (AND for protein complexes, OR for alternative enzymes), there can be several genetic profiles corresponding to a single metabolic profile. We randomly selected 10,000 genetic profiles corresponding to the metabolic profile of T1D and queried each profile programmatically in Enrichr (Chen et al., 2013) through the API. Subsequently, we selected the top five most enriched terms in the KEGG database at p<0.05 for each of the 10,000 gene profiles. Then, we ranked the terms by their occurrence in each profile (Figure S5). An example of the enrichment output of one profile is shown in Table S5. The enrichment analyses of the metabolic reactions in organs and subsystems as groups were carried out through a one-sided hypergeometric test with FDR correction.

### Comparison of flux density estimates

To determine the metabolic effects of insulin in T1D (Figure 4), we first computed the AOS in the T1D model prior to the insulin injection using FVA. We obtained 160,032 solutions (80,160 reactions * 2 (one minimization and one maximization)) in the T1D model AOS, which provided as many numbers of flux values per reaction. Using the empirical flux values per reaction, we estimated the smoothed probability density function of each reaction. Similarly, we collected solutions from the T1D model after the SCIB trial, and we estimated the probability density of the flux values of each reaction. Finally, both density estimates were overlaid on the same axes to compare the effect of insulin on the selection of flux values per enzyme/reaction.

